# NuMA mechanically reinforces the spindle independently of its partner dynein

**DOI:** 10.1101/2024.11.29.622360

**Authors:** Nathan H. Cho, Merve Aslan, Ahmet Yildiz, Sophie Dumont

**Affiliations:** Department of Bioengineering & Therapeutic Sciences, University of California San Francisco, San Francisco, CA, USA; Tetrad Graduate Program, University of California San Francisco, San Francisco, CA, USA; Biophysics Graduate Group, University of California, Berkeley, Berkeley, CA, USA; Department of Physics, University of California, Berkeley, Berkeley, CA, USA; Department of Molecular and Cellular Biology, University of California, Berkeley, Berkeley, CA, USA; Chan Zuckerberg Biohub, San Francisco, CA, USA; Department of Biochemistry & Biophysics, University of California San Francisco, San Francisco, CA, USA

## Abstract

Both motor and non-motor proteins organize microtubules to build the spindle and maintain it against opposing forces. NuMA, a long microtubule binding protein, is essential to spindle structure and function. NuMA recruits the motor dynein to spindle microtubule minus-ends to actively cluster them, but whether NuMA performs other spindle roles remains unknown. Here, we show that NuMA acts independently of dynein to passively reinforce the mammalian spindle. NuMA that cannot bind dynein is sufficient to protect spindle poles against fracture under external force. In contrast, NuMA with a shorter coiled-coil or disrupted self-interactions cannot protect spindle poles, and NuMA turnover differences cannot explain mechanical differences. *In vitro*, NuMA’s C-terminus self-interacts and bundles microtubules without dynein, dependent on residues essential to pole protection *in vivo*. Together, this suggests that NuMA reinforces spindle poles by crosslinking microtubules, using its long coiled-coiled and self-interactions to reach multiple, far-reaching pole microtubules. We propose that NuMA acts as a mechanical “multitasker” targeting contractile motor activity and separately crosslinking microtubules, both functions synergizing to drive spindle mechanical robustness.

## INTRODUCTION

Every cell division, the cell must build the spindle, the microtubule structure that segregates chromosomes. It must also maintain the spindle against forces from both inside and outside the spindle body. Spindle function is essential as failures are linked to cancer and birth defects^1,2^. While we know nearly all molecular-scale players needed for spindle function^3^ how they give rise to the spindle’s robust cellular-scale architecture, dynamics, and mechanics remains poorly understood. Broadly, both motor and non-motor proteins organize microtubules to build and maintain the spindle^4–6^, ultimately giving rise to its function. Motors actively generate force, consuming energy to perform work and move microtubules. In turn, non-motor proteins passively generate force by crosslinking and organizing microtubules without consuming energy, generating friction and elastic forces^7,8^. Spindle proteins are typically thought of as either passive or active force generators, though in reality separating active and passive contributions is often challenging. While there are tools to inhibit active force generators and visualize their action in the spindle, such as chromosome movement, our understanding of passive force generators lags behind due to the relative lack of tools to dissect their contributions.

NuMA (Nuclear Mitotic Apparatus) is a long, complex protein that plays essential roles at different cellular locations and times during the mammalian cell cycle^9^. At mitosis, NuMA is essential for spindle positioning^10–13^ and structure^14,15^. In both roles, NuMA partners with the minus-end-directed motor dynein to actively generate force. In the spindle, NuMA localizes to microtubule minus-ends^16^ and recruits dynein there, driving the clustering of minus-end cargos^17,18^ to build and maintain spindle poles and generate contractile stress in the spindle body^19–22^. Further, recent work shows that NuMA not only recruits dynein to minus-ends, but activates dynein motility there^23,24^. At interphase, NuMA binds DNA and plays a passive, dynein-independent role in nuclear architecture and mechanics^25–28^. Thus, NuMA is an essential, multi-functional protein whose functions must be regulated over space and time.

Whether in mitosis NuMA plays only an active role through dynein or also a passive, dynein-independent role is not known. Purifying full-length NuMA *in vitro* has only been recently achieved^23,24^, and disentangling passive and active roles *in vivo* has been a challenge due to NuMA’s complexity and multiple functions. Through *in vitro* experiments, we know that NuMA binds microtubules^29^, dimerizes^30^, and oligomerizes through self-interactions in a clustering domain^31^, all independent of dynein. In cells, NuMA forms higher order clusters at the cell cortex that recruit dynein to position the spindle^32^ and has recently been proposed to phase separate at the pole^33^. Thus, in principle, NuMA meets all functional requirements to crosslink microtubules on its own: it binds microtubules and can self-interact. As such, NuMA could generate passive force to stabilize the spindle, as has been suggested^14,34^. Notably, NuMA is required for local load-bearing in the mammalian spindle body^35^, suggesting that NuMA helps crosslink spindle microtubules, with or without dynein, and consistently NuMA impacts the rate and synchrony of spindle microtubule transport^36,37^. In some oocytes, NuMA is thought to scaffold microtubule minus-ends at spindle poles, though primarily through dynein rather than on its own^38^. If NuMA did crosslink spindle microtubules on its own, it could in principle synergistically use its active and passive force generation roles, activating dynein at minus-end cargos and crosslinking microtubules, to strengthen the spindle.

Here, we probe NuMA’s passive, dynein-independent contribution to the mechanics of the mammalian mitotic spindle. Using a PDMS-based cell confinement assay, we challenge the spindle with external force and find that NuMA protects human spindle poles from fracturing, independent of any dynein interactions. This establishes a novel, passive role for NuMA in spindle mechanics, independent of its motor partner. We show through different NuMA mutants that NuMA’s long coiled-coil and self-interactions are essential to its ability to mechanically protect spindle poles, and that turnover differences between mutants cannot explain their differential abilities to strengthen the spindle. Finally, we demonstrate that a truncated NuMA can self-interact and bundle microtubules independent of dynein *in vitro*, but that it cannot do so without key residues essential for pole protection *in vivo*. Thus, we propose that NuMA provides mechanical robustness to the spindle by passively crosslinking microtubules, using its long length and ability to form higher-order structures to connect far-reaching, complex microtubule structures at poles. Together with NuMA’s established spindle roles with dynein, this positions NuMA to uniquely multitask in the spindle, playing distinct active and passive roles in maintaining spindle structure. Just like NuMA’s ability to crosslink microtubule cargos could make NuMA/dynein-based force generation more efficient, NuMA/dynein’s ability to remodel the spindle could help NuMA more efficiently and persistently crosslink microtubules.

## RESULTS

### NuMA’s dynein-binding motifs are required for dynein/dynactin recruitment to poles and rescue of NuMA-KO turbulent spindles

To isolate what role, if any, NuMA plays in the spindle without its partner dynein, we sought to generate NuMA mutants that cannot bind dynein. Based on sequence similarities to other dynein-binding proteins and prior work^32,38,39^, NuMA has three proposed dynein-binding motifs: a Hook domain, a CC1-box-like motif, and a Spindly-like motif (Fig. 1A). We designed full-length NuMA mutants with either point mutations in the Spindly-like motif (SpM) to make NuMA-SpM^23,32^ or mutations in both the <u>H</u>ook domain and <u>CC</u>1-box-like motif, previously probed separately^39^, to make NuMA-HCC.

**Figure 1.**
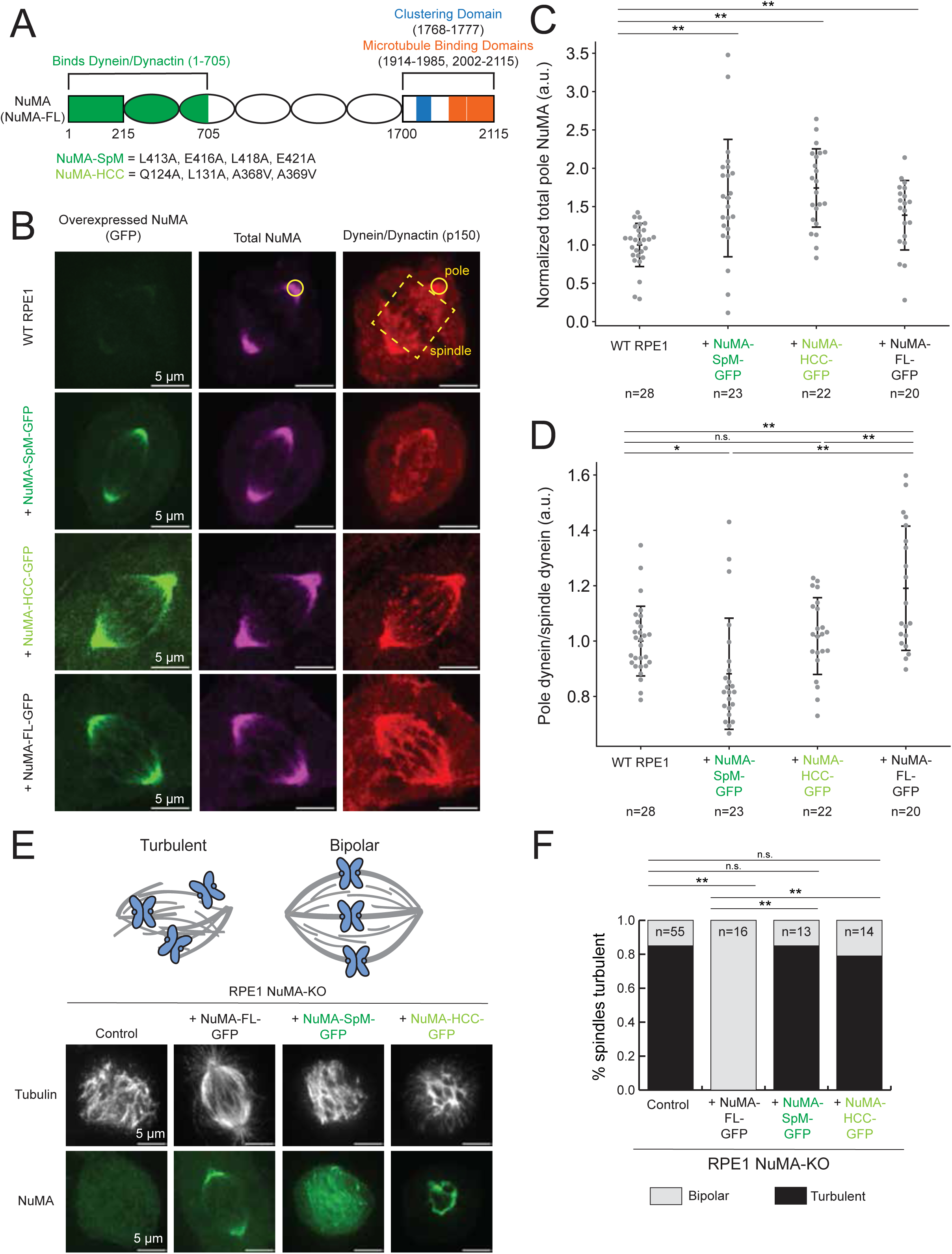
NuMA’s dynein-binding motifs are required for dynein/dynactin recruitment to poles and rescue of NuMA-KO turbulent spindles. (A) Schematic diagram of NuMA and its functional domains, and point mutations used for the NuMA-SpM (from Okumura et al., 2018) and NuMA-HCC mutants (combined individual mutations from Renna et al., 2020) designed to test dynein-independent NuMA functions. (B) Representative confocal images of immunofluorescence staining of WT RPE1 cells and RPE1 cells expressing NuMA-SpM-GFP, NuMA-HCC-GFP, and NuMA-FL-GFP. Cells were stained for GFP (overexpressed NuMA), total NuMA, and dynactin (p150). (C-D) Quantification of total NuMA intensity at poles (C) and ratio of pole dynein/spindle dynein intensities (D), calculated from immunofluorescence images (example of “pole” and “spindle” selections shown on (B)). Values were normalized to mean value for WT RPE1 (control) cells. Data in (C-D) from 2 independent experiments, n=28, 23, 22, and 20 cells. *p < 0.05; **p < 0.005; n.s., not significant; two-sample t-test. Mean ± S.D. (E) Schematic and representative confocal images of immunofluorescence staining of RPE1 NuMA-KO cells as examples of turbulent and bipolar spindle architectures. (F) Percentage of RPE1 NuMA-KO spindles that remain turbulent with exogenous expression of NuMA-FL-GFP, NuMA-SpM-GFP, NuMA-HCC-GFP, or nothing (control). Data from at least 2 independent experiments, n=55, 16, 11, and 12 cells. **p < 0.005; n.s., not significant; Fisher’s Exact Test.

To assess whether NuMA-SpM and NuMA-HCC disrupt NuMA and dynein binding in cells, we took two approaches. First, we examined NuMA and dynein binding by quantifying NuMA and dynein levels at the spindle pole under different conditions (Fig. 1B). Compared to wildtype (WT) RPE1 cells, the total amount of NuMA at spindle poles was increased in RPE1 cell lines stably expressing either NuMA-SpM-GFP, NuMA-HCC-GFP, or full-length NuMA (NuMA-FL-GFP) (Fig. 1C). However, while dynein/dynactin was enriched to the pole relative to control cells in NuMA-FL-GFP expressing cells, dynein/dynactin levels were reduced at the pole in cells expressing NuMA-SpM-GFP (Fig. 1D) and indistinguishable from control in cells expressing NuMA-HCC-GFP despite more NuMA present (Fig. 1D). Therefore, intact dynein-binding NuMA motifs are required for dynein/dynactin recruitment to spindle poles.

Second, we examined the function of these NuMA motifs. Knocking out NuMA or dynein heavy chain leads to a turbulent mammalian spindle phenotype^17^, where the spindle cannot form poles and constantly remodels. This is consistent with NuMA and dynein acting as a complex to cluster microtubule minus-ends into a stable pole^14,15,17^. To test whether NuMA-SpM and NuMA-HCC disrupt NuMA binding to dynein, we performed rescue experiments. In RPE1 cells, we used an inducible CRISPR knockout (KO) approach to delete NuMA^17^, then expressed NuMA-SpM-GFP, NuMA-HCC-GFP, or NuMA-FL-GFP and assessed the extent of rescue of the turbulent spindle phenotype (Fig. 1E). While NuMA-FL-GFP rescued all spindles back to bipolarity, NuMA-SpM-GFP and NuMA-HCC-GFP did not (Fig. 1F). Thus, an intact Spindly-like motif, and either Hook domain or CC1-box-like motif or both, are required to rescue turbulent NuMA-KO spindles, indicating that these domains are required for functional NuMA/dynein complex formation. Given that the mutants do not enrich dynein/dynactin at the spindle pole and do not form a functional NuMA/dynein complex, we consider NuMA-SpM and NuMA-HCC as dynein-binding mutants.

### NuMA provides mechanical robustness to spindle poles independent of dynein

Based on NuMA’s ability to bind microtubules^29^ and self-interact independent of dynein^30,31^, we hypothesized that NuMA can crosslink microtubules and, therefore, provide mechanical robustness to the spindle independent of dynein. Given NuMA’s minus-end localization in cells^17,40,41^, we specifically hypothesized that it passively reinforces spindle poles, maintaining them against force. To test this hypothesis, we confined metaphase cells under a PDMS device to a height of 5 µm to challenge spindles with external force^42–44^ (Fig. 2A) for 45 min (Fig. 2B, Movies S1-S3) and asked if spindle poles could maintain their focused structure. We confirmed that spindle heights before and after 45 min of confinement, and thus the perturbations we applied, were consistent across experiments (Fig. S1A-B). In control WT RPE1 cells, after 45 min of confinement, at least one spindle pole fractured in 35% of spindles (Fig. 2C).

**Figure 2.**
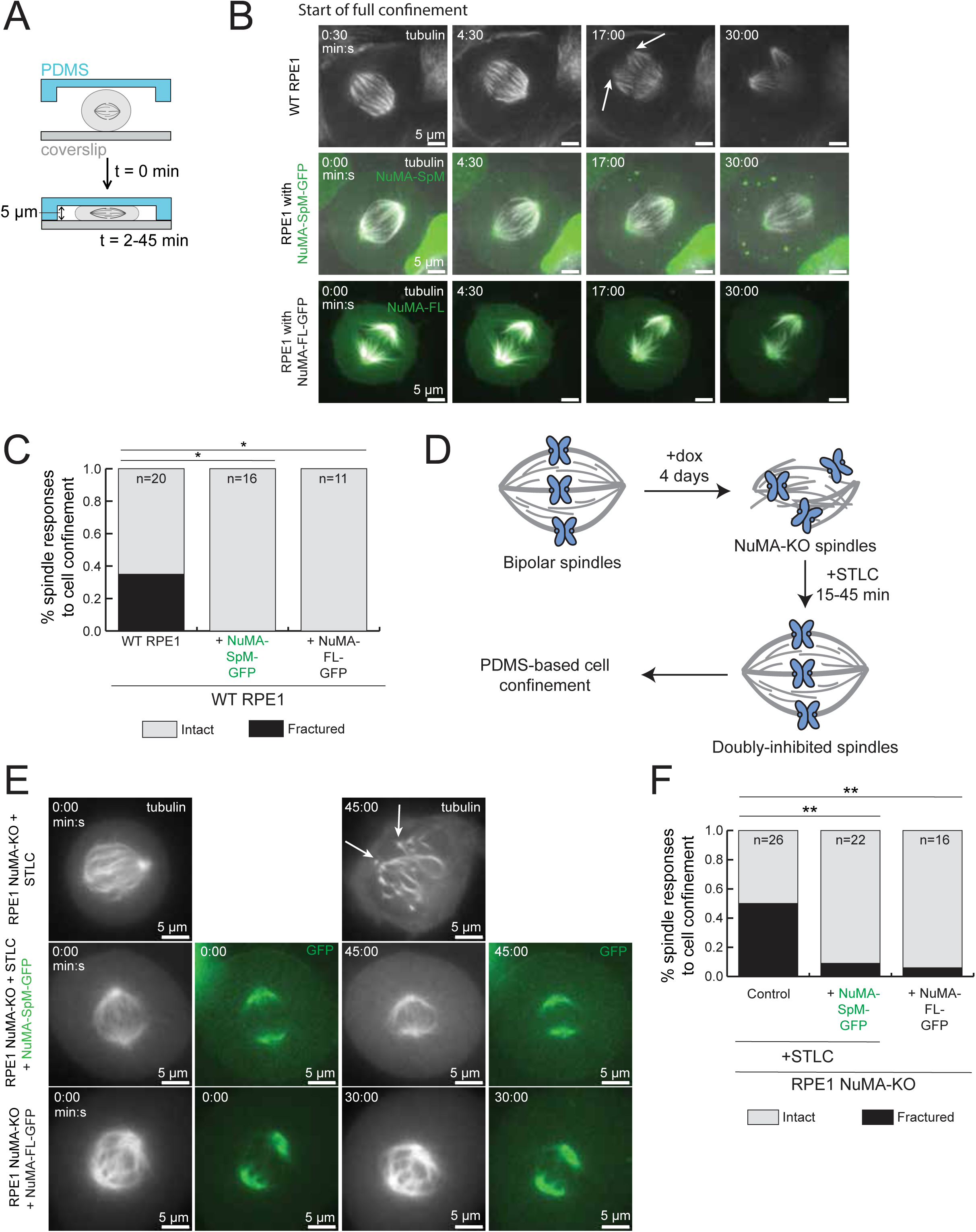
NuMA provides mechanical robustness to spindle poles independent of dynein. **(A)** Schematic illustration of cell confinement experiment to probe spindle mechanical robustness. Confinement to 5 μm height was applied over 2 min, and maintained for 45 min before assessing spindle pole integrity. **(B)** Representative time-lapse confocal images of confinement of a WT RPE1 cell and RPE1 cells overexpressing NuMA-SpM-GFP, NuMA-HCC-GFP, or NuMA-FL-GFP with SiR-tubulin. After cell confinement height reached 5 μm (time = 0:00), spindle pole fracture (white arrows) occurs in the WT RPE1 cell. **(C)** Percentage of RPE1 spindles that fracture upon sustained confinement. n=20, 16, 13, and 11 cells; spindles pooled from ≥3 independent experiments. *p < 0.05; Fisher’s Exact Test. **(D)** Schematic diagram of the workflow for NuMA-KO confinements. Cas9 expression was induced (+dox) for four days to knockout NuMA, resulting in turbulent spindles. Eg5 was then acutely inhibited (STLC) to rescue spindle bipolarity before PDMS-based cell confinement. **(E)** Representative confocal images taken before confinement and after significant confinement time of a RPE1 NuMA-KO + STLC cell, a RPE1 NuMA-KO + STLC + NuMA-SpM-GFP cell, and a RPE1 NuMA-KO + NuMA-FL-GFP cell, all with β-tubulin-HaloTag stained with Janelia Fluor 549 dye. After cell confinement at height 5 µm, spindle pole fracture occurs in some cells, marked by white arrows in the example RPE1 NuMA-KO + STLC cell. **(F)** Percentage of RPE1 NuMA-KO spindles that fracture upon sustained confinement in (E). n=26, 22, and 16 cells; spindles pooled from 4 independent experiments. **p < 0.005; Fisher’s Exact Test.

We first asked whether expression of NuMA-FL-GFP or NuMA-SpM-GFP (Fig. 2A) on top of endogenous NuMA was sufficient to reduce the spindle pole fracture rate under confinement. Here and forward, we chose NuMA-SpM over NuMA-HCC as it appeared more efficient (Fig. 1D and 1F) at inhibiting dynein binding and function. Importantly, we expressed NuMA-SpM-GFP and NuMA-FL-GFP at similar levels, as assessed by fluorescent intensity measurements at spindle poles. Overexpressing NuMA-FL leads to more NuMA at poles (Fig. 1C) and reduces the incidence of spindle pole fracture under confinement (Fig. 2C), demonstrating that NuMA-FL can mechanically reinforce spindle poles. If NuMA’s ability to mechanically reinforce poles is dynein-independent, then expressing NuMA-SpM on top of endogenous should reduce spindle pole fracture rate similar to NuMA-FL, and this is what we found (Fig. 2C). In fact, expression of NuMA-SpM and NuMA-FL on top of endogenous NuMA each completely eliminated spindle pole fracture under confinement (Fig. 2C). Therefore, NuMA-SpM, which cannot interact with dynein, is as competent as NuMA-FL at mechanically protecting spindle poles, when expressed on top of endogenous NuMA.

In principle, NuMA-SpM’s ability to protect spindle poles (Fig. 2C) could come from its interaction with endogenous NuMA which can interact with dynein. To test this possibility, we performed the same assay in inducible NuMA-KO RPE1 cells. Because NuMA-KO spindles are turbulent^17^, we inhibited Eg5 with S-trityl-L-cysteine (STLC), NuMA/dynein’s opposite motor^45,46^, to make non-turbulent bipolar spindles without NuMA before beginning cell confinement (Fig. 2D). For the NuMA-FL-GFP condition, STLC was not added since NuMA-FL-GFP expression restores bipolarity on its own (Fig. 1E) and STLC addition can lead to monopoles^47^. Previous work has shown that doubly-inhibited (NuMA-KO + STLC) spindles are less mechanically robust than spindles in WT RPE1 cells^48^. In agreement, we find that doubly-inhibited control spindles exhibit pole fracture 50% of the time (Fig. 2E-F). Expressing NuMA-SpM-GFP or NuMA-FL-GFP in this NuMA-KO background significantly decreased the spindle pole fracture rate compared to control (Fig. 2F), similar to spindles with endogenous NuMA (Fig. 2C) though now with more variable spindle shapes. The time to spindle fracture is similar between doubly-inhibited control, NuMA-SpM-GFP, and NuMA-FL-GFP (Fig. S1C), suggesting that NuMA expression impacts the likelihood of fracture as opposed to the kinetics of fracture. Further, given that NuMA-KO spindles do not have endogenous NuMA interacting with astral microtubules at the cell cortex, these findings are inconsistent with cortical NuMA being responsible for differences in spindle fracture between conditions. Taken together, we conclude that NuMA mechanically protects spindle poles independent of any NuMA interactions with dynein. Thus, NuMA plays a passive, dynein-independent role in maintaining metaphase spindle structure.

### NuMA’s coiled-coil and clustering domain are both required for spindle pole protection

We next asked how NuMA provides mechanical robustness to spindle poles without dynein. In principle, NuMA could passively reinforce the spindle through crosslinking microtubules by binding microtubules^29^, forming a homodimer through its coiled-coil^30^ and assembling into higher order structures through a C-terminus clustering domain^31,32^. Further, we hypothesized that NuMA’s unusually long length (∼210 nm), which mostly comes from its coiled-coil^30,31^, increases the distance over which it can bind microtubules and thus enhances its ability to crosslink microtubule structures. To test whether the coiled-coil and clustering domain are required for spindle pole protection, we generated combination NuMA mutants (Fig. 3A) that cannot bind dynein and which either have a shortened coiled-coil (NuMA-SpM-Bonsai), cannot self-interact through the clustering domain (NuMA-SpM-5A-3), or cannot dimerize or self-interact through the clustering domain (NuMA-Cterm-5A-3). Consistent with the 5A-3 mutation disrupting the formation of higher order NuMA structures, it disrupts oligomerization at the metaphase cortex^32^, and we found that NuMA-Cterm-5A-3-GFP disrupts the formation of nuclear NuMA filaments that form when we express NuMA-Cterm-GFP in RPE1 cells (Fig. 3B). These filaments, resembling previously reported nuclear NuMA filaments^27,28^, occur in 35% of nuclei and co-localize with tubulin (Fig. 3B), resist nocodazole treatment (Fig. S2A), and can organize endogenous NuMA-FL and even DNA (Fig. S2B).

**Figure 3.**
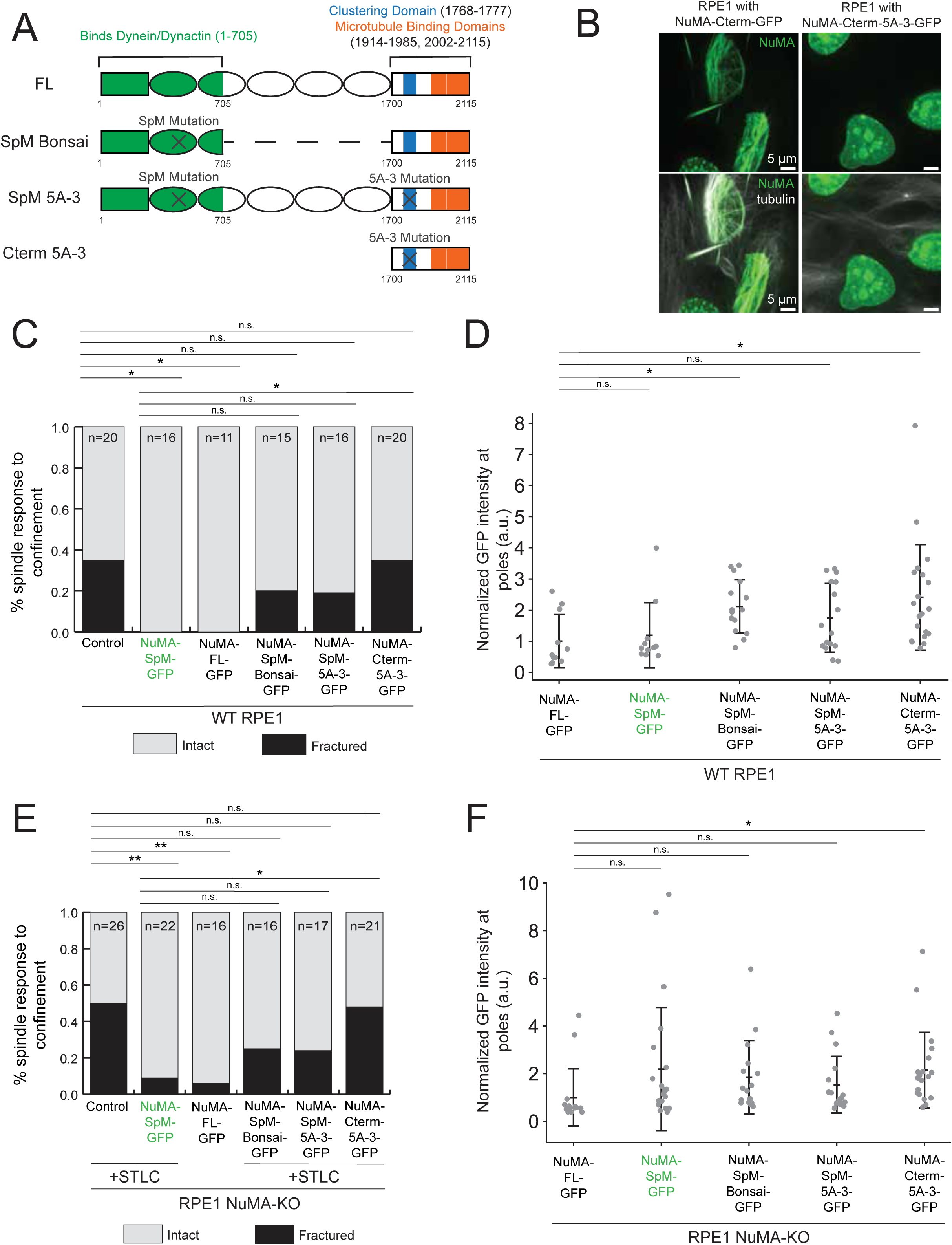
NuMA coiled-coil and clustering domain are both required for spindle pole protection. **(A)** Schematic maps of different NuMA mutants used. ‘SpM-Bonsai’ has the SpM mutation (Fig. 1A) and a truncation of the coiled-coil. ‘SpM-5A-3’ has the SpM mutation and 5A-3 mutation (Okumura et al., 2018) which disrupts NuMA self-binding. ‘Cterm-5A-3’ is a truncation with the 5A-3 mutation. **(B)** Representative confocal images of interphase RPE1 cells overexpressing NuMA-Cterm-GFP or NuMA-Cterm-5A-3-GFP with SiR-tubulin, indicating that 5A-3 mutation disrupts filament formation and self-interactions. **(C)** Percentage of RPE1 spindles that fracture upon sustained confinement, including data from Fig. 2C. n=20, 16, 13, 11, 15, 16, and 20 cells; spindles pooled from ≥3 independent experiments. *p < 0.05; n.s., not significant; Fisher’s Exact Test. **(D)** NuMA-GFP expression at spindle poles in RPE1 cells expressing various NuMA-GFP constructs (Fig. 2C and Fig. 3C), normalized to mean value of NuMA-FL-GFP. From the same cells as (C). *p < 0.05; n.s., not significant; two-sample t-test. Mean ± S.D. **(E)** Percentage of RPE1 NuMA-KO spindles that fracture upon sustained confinement, including data from Fig. 2F. n=26, 22, 16, 16, 17, and 21 cells; spindles pooled from ≥3 independent experiments. *p<0.05; **p < 0.005; n.s., not significant; Fisher’s Exact Test. **(F)** NuMA-GFP expression at spindle poles in RPE1 NuMA-KO cells expressing various NuMA-GFP constructs (Fig. 2F and Fig. 3F), normalized to mean value of NuMA-FL-GFP. From the same cells as (E). *p < 0.05; n.s., not significant; two-sample t-test. Mean ± S.D.

To test the contributions of NuMA’s coiled-coil and clustering domain to passive spindle pole reinforcement, we confined RPE1 cells stably expressing NuMA-SpM-Bonsai-GFP, NuMA-SpM-5A-3-GFP, or NuMA-Cterm-5A-3-GFP as above (Fig. 2). We found that expressing NuMA-SpM-Bonsai-GFP or NuMA-SpM-5A-3-GFP did not eliminate spindle pole fracturing as NuMA-SpM-GFP and NuMA-FL-GFP had done, but decreased pole fracturing compared to control WT cells (Fig. 3C, Fig. S3A). This suggests that NuMA’s length and self-interactions both contribute to passive reinforcement of spindle poles. NuMA-Cterm-5A-3, however, did not decrease spindle pole fracturing compared to control (Fig. 3C, Fig. S3A), consistent with NuMA’s coiled-coil and clustering domain both being necessary for spindle pole reinforcement. While NuMA-SpM-5A-3-GFP and NuMA-Cterm-5A-3-GFP sometimes mis-localize to chromosomes (Fig. S3A), these NuMA constructs were at a similar (NuMA-SpM-5A-3-GFP) or higher (NuMA-SpM-Bonsai-GFP and NuMA-Cterm-5A-3-GFP) intensity level as NuMA-SpM-GFP and NuMA-FL-GFP at spindle poles (Fig. 3D). Therefore, differences in response to confinement cannot be explained by reduced NuMA localization to the pole.

In principle, here too endogenous NuMA could interact with NuMA-SpM-Bonsai and NuMA-SpM-5A-3 and confound interpretations. To test this idea, we repeated the same experiments in NuMA-KO cells and found the same things as when endogenous NuMA were present: NuMA-KO spindles with NuMA-Cterm-5A-3-GFP fractured indistinguishably from control NuMA-KO spindles (Fig. 3E, Fig. S3B), and NuMA-KO spindles with NuMA-SpM-Bonsai-GFP or NuMA-SpM-5A-3-GFP had decreased spindle pole fracturing relative to control, but did not eliminate spindle pole fracturing as NuMA-SpM-GFP and NuMA-FL-GFP did (Fig. 3E, Fig. S3B). Again, and importantly, NuMA levels at spindle poles were similar (NuMA-SpM-Bonsai-GFP and NuMA-SpM-5A-3-GFP) or higher (NuMA-Cterm-5A-3-GFP) than NuMA-SpM-GFP and NuMA-FL-GFP, and thus differences in response to force are not due to reduced NuMA levels at poles (Fig. 3F). Together, these findings indicate that NuMA’s coiled-coil and clustering domain both contribute to NuMA’s passive, dynein-independent spindle pole protection. We propose that the clustering domain is essential to crosslink microtubules by linking two or more separate microtubules and that a long coiled-coil helps crosslink distant microtubules and mechanically protect the pole.

### Spindly-like motif, coiled-coil, and clustering domain each impact NuMA turnover at spindle poles but cannot explain differences in pole protection

Differences in the ability of NuMA constructs to protect spindle poles against fracture (Fig. 2-3) could either stem from changes in the turnover rate of NuMA at poles or from changes in NuMA crosslinks microtubules at poles. To test these models, we measured the turnover rate of NuMA-FL-GFP^49^ and NuMA-GFP mutants at spindle poles using <u>F</u>luorescence <u>R</u>ecovery <u>A</u>fter <u>P</u>hotobleaching (FRAP, Fig. 4A, Movie S4).

**Figure 4.**
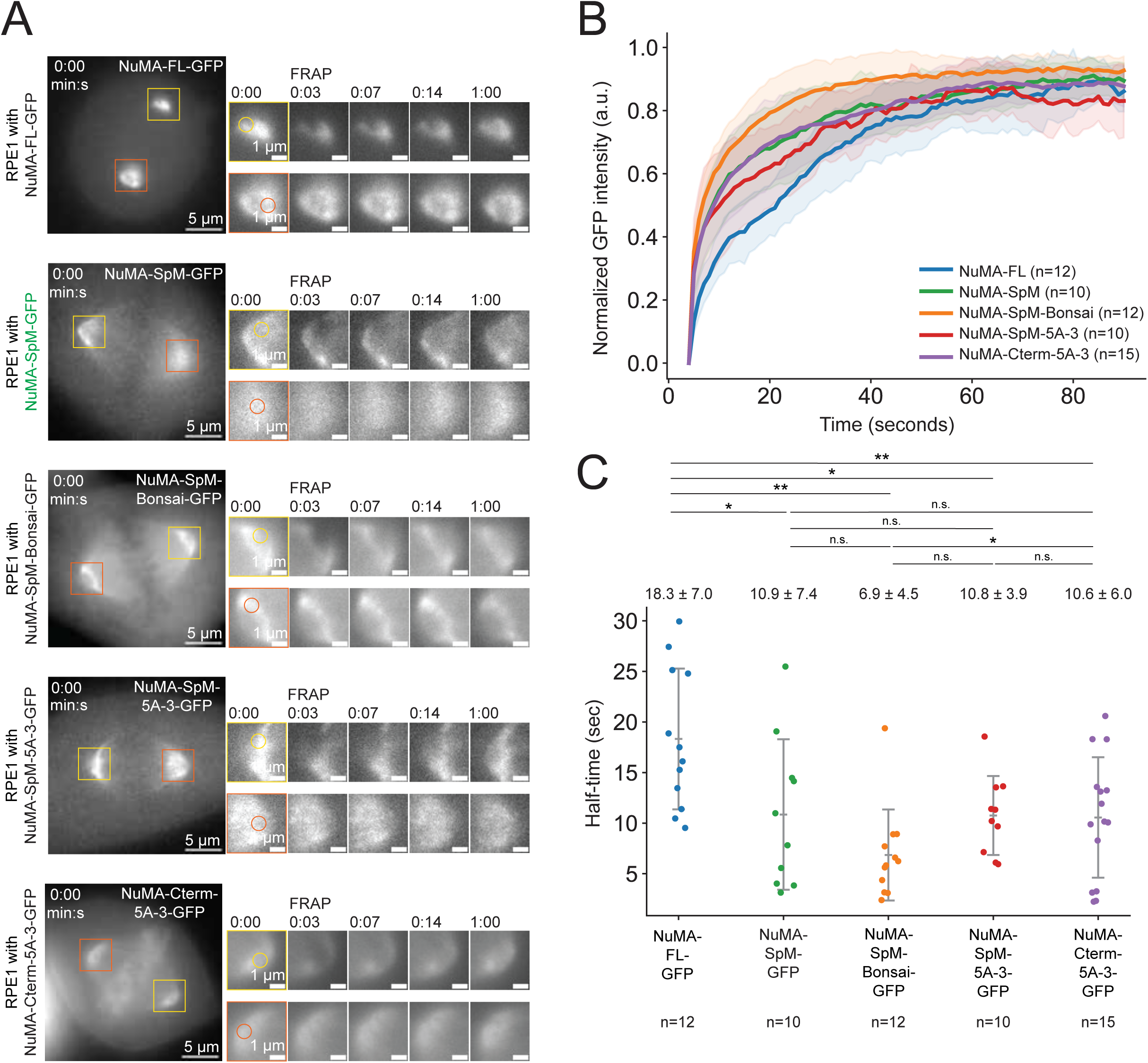
Spindly-like motif, coiled-coil, and clustering domain each impact NuMA turnover at the pole, but cannot explain differences in spindle pole protection. **(A)** Representative time-lapse widefield images and zoom insets (yellow/orange squares) of spindle poles of FRAP on a RPE1 cell expressing NuMA-FL-GFP, NuMA-SpM-GFP, NuMA-SpM-Bonsai-GFP, NuMA-SpM-5A-3-GFP, or NuMA-Cterm-5A-3-GFP. Spot (yellow circle position) is bleached between t = 0:02-0:03, and intensity compared to spot in non-targeted pole (orange circle position). **(B)** FRAP GFP intensity (normalized to other, non-targeted pole) of different NuMA-GFP constructs in metaphase RPE1 cells. n=12, 10, 12, 10, and 15 cells; spindles pooled from at least 2 independent experiments. **(C)** Total half-time of recovery for different NuMA-GFP constructs from Fig. 4B. *p < 0.05; **p<0.005; n.s., not significant; two-sample t-test. Mean ± S.D.

First, we measured turnover of NuMA-FL-GFP and NuMA-SpM-GFP to assess if NuMA’s dynein association changes its turnover dynamics (Fig. 4B). NuMA-SpM-GFP had a shorter recovery half-time at the pole than NuMA-FL-GFP (10.9 ± 7.4 s vs. 18.3 ± 7.0 s, Fig. 4C). Thus, dynein-independent NuMA turns over more quickly than NuMA-FL which can interact with dynein, consistent with NuMA binding the pole through both dynein-dependent and -independent mechanisms. Second, we measured turnover of NuMA-SpM-Bonsai-GFP, NuMA-SpM-5A-3-GFP, and NuMA-Cterm-5A-3-GFP (Fig. 4B). NuMA-SpM-5A-3-GFP had a similar recovery half-time at the pole to NuMA-SpM-GFP (10.8 ± 3.9 s vs. 10.9 ± 7.4 s, Fig. 4C), suggesting that loss of self-interactions does not have a significant effect on turnover at the pole. NuMA-SpM-Bonsai-GFP had a shorter half-time of recovery at the pole compared to NuMA-SpM-GFP (6.9 ± 4.5 s vs. 10.9 ± 7.4 s, Fig. 4C), and thus reducing NuMA’s length increases its turnover, consistent with a longer NuMA molecule having a higher chance of being microtubule-bound.

Overall, differences in the turnover of NuMA constructs cannot explain differences in the mechanical protection of spindle poles. NuMA-SpM and NuMA-FL have different turnover rates (Fig. 4B-C) yet are similarly able to protect spindle poles with (Fig. 2C) and without (Fig. 2F) endogenous NuMA. Similarly, NuMA-SpM-Bonsai and NuMA-SpM-5A-3 have noticeably different turnover rates (Fig. 4B-C) but provide similar mechanical protection mechanics both with (Fig. 3C) and without (Fig. 3E) endogenous NuMA. Lastly, NuMA-Cterm-5A-3 has a similar turnover rate to NuMA-SpM (Fig. 4B-C) but does not protect spindle poles, unlike NuMA-SpM with (Fig. 3C) and without (Fig. 3E) endogenous NuMA. Thus, the loss of spindle pole robustness with NuMA-SpM-Bonsai, NuMA-SpM-5A-3, and NuMA-Cterm-5A-3 cannot be attributed to reduced or increased turnover of NuMA at the pole. These findings support the model that NuMA’s passive role in spindle pole robustness results from NuMA crosslinking microtubules at the pole.

### NuMA C-terminus can self-interact and bundle microtubules *in vitro* without dynein and does so more effectively than NuMA C-terminus 5A-3

Our data support the model that NuMA self-interacts and crosslinks microtubules to reinforce spindle poles independent of dynein. Interpretation of key data supporting this model requires that the NuMA 5A-3 mutation (Fig. 3) indeed disrupts self-interactions to disrupt crosslinking. Further, this model requires that NuMA is sufficient to crosslink microtubules *in vitro*, which has previously only been shown with portions of *Xenopus* NuMA^14^ or a synthetic dimer NuMA^34^. We tested both requirements *in vitro* by expressing and purifying (Fig. 5A) human NuMA-Cterm-mNeonGreen (1700-2115) and NuMA-Cterm-5A-3-mNeonGreen (as in Fig. 3 and 4). Using mass photometry, we found that while NuMA-Cterm often exists as a monomer (Fig. 5B-C), it also exists as a dimer (Fig. 5B-C) and high-order assemblies of molecules (Fig. 5B-C). Conversely, NuMA-Cterm-5A-3 exists more dominantly as a monomer (Fig. 5B-C), consistent with the 5A-3 mutations disrupting NuMA’s self-interactions.

**Figure 5.**
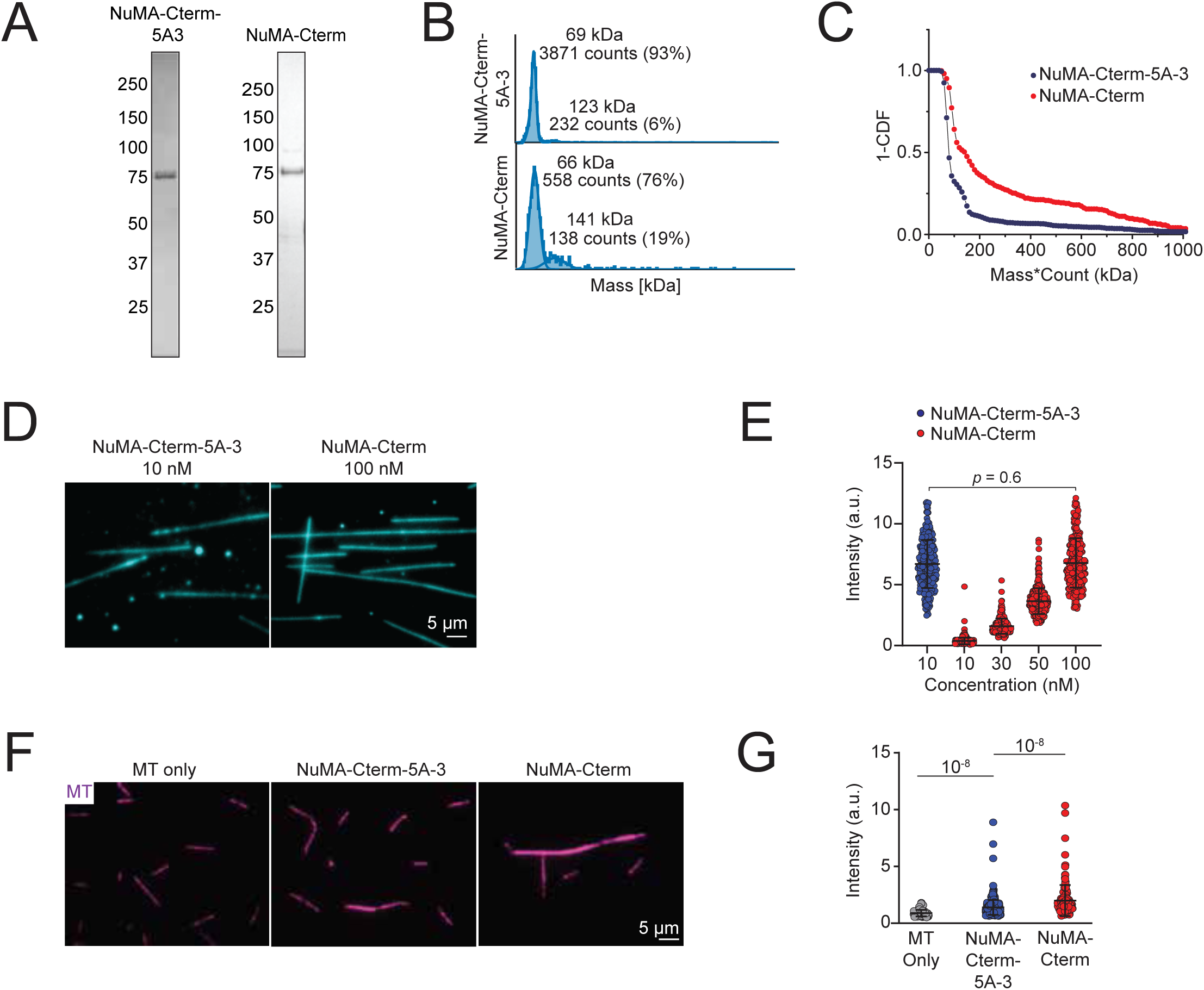
NuMA C-terminus can self-interact and bundle microtubules *in vitro* without dynein and does so more effectively than NuMA C-terminus 5A-3. (A) Denaturing gel images of NuMA-Cterm-GFP and NuMA-Cterm-5A-3-GFP after gel filtration. The numbers on the left represent molecular weight in kDa. **(B)** Mass photometry traces of NuMA-Cterm-GFP and NuMA-Cterm-5A-3-GFP (each 20 nM). **(C)** 1 – cumulative distribution (CDF) of mass x count plot of mass photometry measurements for NuMA-Cterm-GFP and NuMA-Cterm-5A-3-GFP. **(D)** Representative images of 10 nM NuMA-Cterm-5A-3-mNeonGreen and 100 nM NuMA-Cterm-mNeonGreen on surface immobilized microtubules. **(E)** Fluorescence intensity analysis of NuMA proteins per length of a microtubule in each condition. Mean ± S.D. For data in (D-E), n=270, 561, 298, 357, and 275 microtubules; microtubules pooled from at least 2 independent experiments. *P* values were calculated with student’s t-test. **(F)** Representative TIRF images of Cy3 labelled microtubules in the presence of nothing (MT only), NuMA-Cterm-5A-3-GFP, or NuMA-Cterm-GFP in a microtubule bundling assay. **(G)** Fluorescence intensity of tubulin in microtubule bundling assay. Mean ± S.D. For data in (F-G), n=92, 582, 150 microtubules; microtubules pooled from at least 2 independent experiments. *P* values were calculated with student’s t-test.

To test NuMA’s ability to bind and crosslink microtubules, and the impact of 5A-3 mutations, we used TIRF microscopy in microtubule binding and bundling assays. We found that NuMA-Cterm-5A-3 has a higher microtubule affinity than NuMA-Cterm, which may be due to the clustering of NuMA negatively regulating its microtubule binding (Fig. 5D). Under similar microtubule binding conditions, NuMA-Cterm can bundle microtubules independent of dynein, while NuMA-Cterm-5A-3 is much less efficient at doing so (Fig. 5E-F), consistent with NuMA-Cterm-5A-3 not forming higher order NuMA structures in the nucleus as NuMA-Cterm did (Fig. 3B), and with NuMA-Cterm-5A-3’s inability (and NuMA-SpM-5A-3’s reduced ability compared to NuMA-SpM) to protect spindle poles (Fig. 3C-F). Overall, these findings show that NuMA-Cterm can self-interact and bundle microtubules, both independent of dynein and despite reduced microtubule affinity, and that the 5A-3 residues essential for pole protection in cells are also essential for self-interaction and bundling *in vitro*. More broadly, this *in vitro* work supports the model that NuMA can passively crosslink microtubules and thereby reinforce spindle poles without its motor partner.

## DISCUSSION

NuMA is essential for spindle structure and function, and it has long been known to bind and work with the motor dynein to actively build the spindle. In contrast, here we show that NuMA can mechanically reinforce and maintain spindle poles through passive force generation, independent of dynein and its energy consumption. We demonstrate that NuMA can protect spindle poles against fracture without binding dynein (Fig. 1-2), that this mechanical role requires both NuMA’s long coiled-coil and self-interaction (Fig. 3), and cannot simply be explained by differences in NuMA turnover kinetics (Fig. 4). Finally, we show that NuMA-Cterm can self-interact and bundle microtubules *in vitro*, and that NuMA-C-term-5A-3 cannot (Fig. 5), mirroring residues essential for spindle pole protection (Fig. 3). Our findings support a model where NuMA forms higher order structures that crosslink far-reaching microtubules to protect spindle poles (Fig. 6). We propose that NuMA is a mechanical “multitasker” in the spindle, performing active and passive roles that are distinct and reinforce each other to make the spindle mechanically robust.

**Figure 6.**
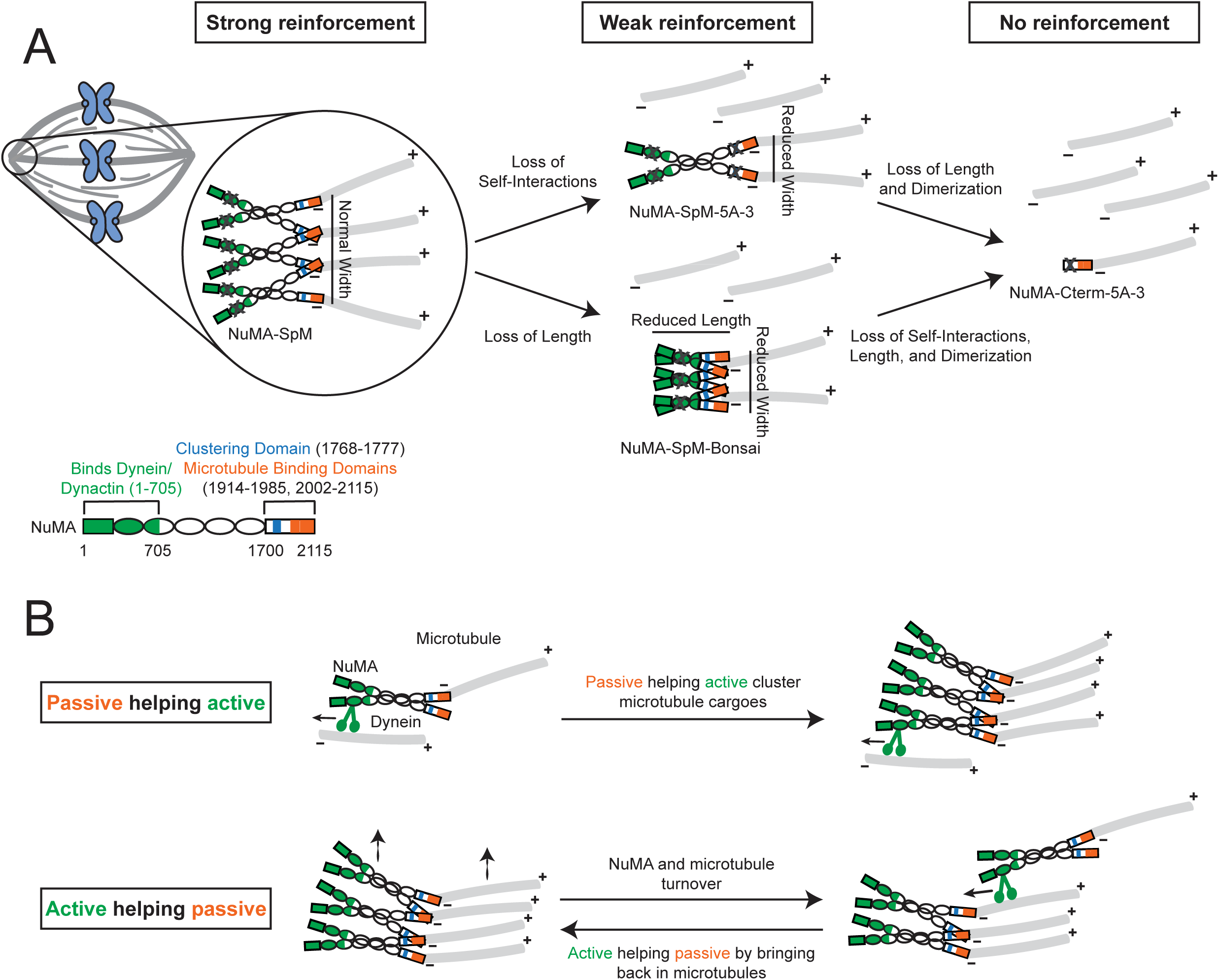
Model for how NuMA passively crosslinks microtubules and synergistically targets dynein force to protect the spindle. **(A)** Model for how NuMA (bottom left) passively crosslinks (orange) spindle microtubules (grey) without its partner dynein (SpM mutation, grey X on green) using its ability to self-interact through its C-terminus (top, blue) and its long coiled-coil (bottom, white). Loss of either (NuMA-SpM-5A-3 (grey X on blue) or NuMA-SpM-Bonsai (shorter white coiled-coil), respectively) or both (NuMA-Cterm-5A-3, right) reduces NuMA’s ability to mechanically reinforce the spindle. Self-interactions and a long coiled-coil could, for example, boost effective crosslinking by increasing complexity or flexibility, respectively, of a NuMA cluster. NuMA’s long and self-interacting structure helps it crosslink microtubules over large distances and complex structures, such as spindle poles (far left). **(B)** Model for NuMA as a mechanical “multitasker” with passive (crosslinking, orange) and active (dynein/NuMA-based minus-end transport, green) mechanical roles, with both roles synergizing to maintain spindle structure. NuMA’s passive role helps its active role by crosslinking microtubule (grey) cargoes, making dynein/NuMA-based active transport more efficient (top). Conversely, NuMA’s active role helps restore its passive crosslinking role by bringing NuMA molecules and microtubules back into a cluster after they turnover (bottom).

Our findings support a model where NuMA can passively crosslink spindle microtubules to reinforce spindle poles. Previous *in vivo* work suggested that NuMA could crosslink microtubules, though it was not clear whether this was through dynein or not^14,35,38,50^, and *in vitro* work showed that portions of *Xenopus* NuMA’s C-terminus^14^ and synthetic dimer NuMA constructs^34^ can bundle microtubules. Here, we show that human NuMA’s C-terminus can bundle microtubules *in vitro* dependent on key residues (Fig. 5C-D), that NuMA’s full-length coiled-coil and self-interactions through these residues are necessary for its passive crosslinking role in the spindle (Fig. 3), and that this role is independent of dynein (Fig. 2). Whether this role is independent of other pole focusing proteins, such as HSET^38^, remains an open question. Both a full-length coiled coil domain and ability to self-interact can in principle increase the distance over which NuMA can bind and crosslink microtubules, and increase the size and complexity of NuMA-microtubule bundles (Fig. 6A). Notably, while NuMA’s long length is dispensable for its active dynein-based role in spindle assembly^17^, here we show that NuMA’s length is essential for spindle pole maintenance (Fig. 3). This suggests a function for NuMA’s long (∼1500 amino acid, ∼210 nm) coiled-coil in the spindle, a long-standing question. Systematically tuning NuMA’s length and self-interaction stoichiometry and measuring its ability to reinforce spindle poles could ultimately reveal how NuMA crosslinks pole microtubules, and the architecture of the pole. This could also reveal whether different NuMA functions regulate each other, for example NuMA clustering negatively regulating microtubule binding as suggested by our *in vitro* work (Fig. 5D). Similarly, understanding how NuMA’s length and self-interactions give rise to its nuclear function^27,51,52^ will help uncover how it supports nuclear formation and mechanics.

NuMA’s active and passive roles – set within the same protein – are well suited to synergistically support each other (Fig. 6B). NuMA’s passive role in reinforcing microtubule structures can in principle not only stabilize spindle poles, but crosslink microtubule cargoes as NuMA activates dynein^23,24^ and dynein/NuMA transport microtubules to spindle poles^14^, making this process more efficient. Conversely, dynein/NuMA’s active transport of microtubules can help maintain crosslinking by continuously bringing microtubules minus-ends together for NuMA to crosslink, for example after microtubule minus-ends drift away as NuMA turns over on the order of seconds (Fig. 4). In both directions, NuMA’s distinct active and passive force generation roles support each other and make for an elegant and efficient usage of a large protein during mitosis. Consistent with this, recent work has showed that NuMA/dynein’s active role alone is not sufficient for aster formation *in vitro*^23^. Instead, efficient aster formation requires full-length mitotically phosphorylated NuMA^23^, which suggests the active and passive roles are both needed. To our knowledge, this makes NuMA biophysically unique amongst currently understood spindle molecules. For example, while spindle motors such as Eg5^53^, HSET^54^, and even dynein itself^55,56^ can crosslink microtubules while transporting them, this crosslinking is a byproduct of the motor’s active role, not a distinct passive role. Similarly, spindle microtubule crosslinkers such as PRC1^57–59^ can recruit motors^60^, but do not directly activate the motor as NuMA does with dynein. Also consistent with synergy between active and passive roles, modeling work on actomyosin systems^61^ predicts that an active contractile system (such as the spindle pole) will have maximal contractility with both an end-to-end crosslinker (like NuMA) and a motor (like NuMA/dynein). To our knowledge, NuMA is the only dynein activating adaptor with microtubules as cargo, and also the only to form higher order self-assemblies. Looking forward, it will be important to determine whether a single NuMA molecule multi-tasks at once or switches between passive and active roles, and what regulates this switch if it exists.

Our work puts forward the question of whether NuMA’s phosphorylation impacts its active or passive roles, and the relative contribution of each role. NuMA’s phosphorylation regulates its localization and microtubule binding affinity during mitosis, and changes at mitotic entry, anaphase entry, and mitotic exit^40,41,62,63^. Phosphorylation is key to regulating NuMA’s functions over time through the cell cycle and over space at different locations, for example at spindle poles and the cell cortex during mitosis. We know that NuMA’s phosphorylation can impact its affinity to dynein and thus its active role^23^. Given that NuMA’s C-terminus is phosphorylated at its clustering motif and microtubule binding sites^40,41,62,63^, we propose that NuMA’s phosphorylation also regulate its passive role by changing its self-interactions and microtubule affinity. Such regulation may also allow NuMA to play related but structurally distinct passive roles in the spindle and nucleus.

Together, our work raises the question of whether spindle molecules that can generate force both actively and passively, in distinct roles with and without energy consumption, can play unique biophysical roles in cells that others cannot. The framework of collective phenomena and simplified *in vitro* and computational systems^61,64^ will be critical for answering this question, allowing us to test necessity and sufficiency of NuMA’s active and passive roles, and their combination. Determining whether and how NuMA’s passive and active roles together give rise to a new function that is more than sum of its roles will be an important future direction.

## ACKNOWLEDGMENTS

We thank Lila Neahring, Christina Hueschen, Andrea Serra-Marques, Matt Thomson, and members of the Dumont and Yildiz Labs for helpful discussions, and members of the Dumont Lab for critical reading of the manuscript. We thank Zhidong Tan for photomarking help. FRAP experiments were performed at the UCSF Center for Advanced Light Microscopy with assistance from Kari Herrington, on an OMX-SR obtained using funding from NIH 5R35GM118119, the UCSF Program in Breakthrough Biomedical Research funded in part by the Sandler Foundation, the UCSF Research Resource Fund Award, and the Howard Hughes Medical Institute. This work was supported by an AHA Predoctoral Fellowship (908941), UCSF Discovery Fellowship, and NIH F31CA275394 (N.H.C.), NIH GM136414 and NSF MCB-1055017 and MCB-1617028 (A.Y.), and NIH R35GM136420 and NSF 1548297 Center for Cellular Construction (S.D.). S.D. is a Chan Zuckerberg Biohub Investigator.

## AUTHOR CONTRIBUTIONS

Conceptualization, N.H.C., A.Y., and S.D.; methodology, N.H.C. and M.A.; software, N.H.C. and M.A.; validation, N.H.C. and M.A.; formal analysis, N.H.C. and M.A.; investigation, N.H.C. and M.A.; resources, N.H.C., M.A., A.Y., and S.D.; data curation, N.H.C. and M.A.; writing – original draft, N.H.C.; writing – review and editing, N.H.C., M.A., A.Y., and S.D.; visualization, N.H.C. and M.A.; supervision, A.Y. and S.D.; funding acquisition, N.H.C., A.Y., and S.D.

## METHODS

### Cell culture (Fig. 1-4)

hTERT-RPE1 cells (female human retinal epithelial cells) were purchased from ATCC (CRL-4000). Tet-on inducible CRISPR/Cas9 NuMA-KO RPE1 cells^17^ additionally expressing β-tubulin-HaloTag^65^ were from prior work. All RPE1 cells were grown in DMEM/F12 (11320; Thermo Fisher Scientific) supplemented with 10% tetracycline-screened FBS (PS-FB2; Peak Serum). SpCas9 expression was induced by adding 1 µg/mL doxycycline hyclate (D9891; Sigma-Aldrich) 4 days before each experiment and refreshed at 24 and 48 hours. All cells were maintained at 37 °C and 5% CO_2_.

### Lentiviral plasmids and cell line construction (Fig. 1-4)

The coding sequence of all NuMA-GFP constructs (NuMA-FL, NuMA-SpM, NuMA-HCC, NuMA-SpM-Bonsai, NuMA-SpM-5A-3, NuMA-Cterm, and NuMA-Cterm-5A-3) were cloned into a puromycin-resistant lentiviral vector (Addgene #114021). Lentivirus for each construct was produced in HEK293T cells that were purchased from ATCC (CRL-3216) and grown in DMEM/F12 supplemented with 10% tetracycline-screened FBS. RPE1 cells were either transiently infected with these lentivirus (Fig. 1E-F) or infected with lentivirus and either selected using 5 µg/mL puromycin (WT RPE1 background cell lines, Fig. 2C, 3C, 3D, and 4) or FACS sorted (NuMA-KO cell lines, Fig. 2F, 3E, and 3F) to generate stable polyclonal cell lines.

### Dyes and drug treatments (Fig. 2-4)

To visualize microtubules, WT RPE1 cells were treated with 100 nM SiR-tubulin and 10 µM verapamil for 60 min prior to imaging (CY-SC002; Cytoskeleton, Inc.) (Fig. 2B, 2C, 3B, 3C, 4A, S2A, and S3A). Alternatively, inducible NuMA-KO RPE1 cells, which stably express β-tubulin-HaloTag, were treated with 100 nM Janelia Fluor 549 (GA1110; Promega) for 15 min prior to imaging (Fig. 2E, 2F, 3E, and S3B).

For experiments with doubly-inhibited RPE1 cells (Fig. 2F, 3E, and 3F), STLC (164739; Sigma-Aldrich) was added to a final concentration of 5 µM for 15-45 minutes before confinement, as assessed by restoration of spindle bipolarity. To depolymerize microtubules, nocodazole (M1404; Sigma-Aldrich) was added to a final concentration of 5 µM for 15-30 min prior to imaging (Fig. S2A). Given the lack of interphase cytoplasmic microtubules, we determined this was sufficient nocodazole treatment.

### Immunofluorescence (Fig. 1)

Cells were plated on acid-cleaned, poly-*L*-lysine coated, #1.5 25 mm coverslips for 1–3 days. Coverslips were washed in phosphate-buffered saline (PBS), fixed in pre-chilled 100% MeOH pre-chilled to −20 °C for 3-5 min, and washed again in PBS. Coverslips were blocked in TBST (0.05% Triton X-100 in tris-buffered saline) containing 2% (wt/vol) bovine serum albumin (BSA). Antibodies were diluted in TBST containing 2% BSA and incubated for 1 hour (primary antibodies) or 45 minutes (secondary antibodies) at room temperature, followed by four washes in TBST. DNA was labeled with 1 µg/mL Hoechst 3342 prior to mounting on slides with ProLong Gold Antifade Mountant (P36934; Thermo Fisher Scientific). The following primary antibodies were used: rabbit anti-NuMA (1:300, NB500-174, RRID: AB_10002562; Novus Biologicals), mouse anti-p150 [Glued] (1:400, 610474, RRID: AB_397846; BD Biosciences), mouse anti-α-tubulin (1:1000, T6199, RRID: AB_477583; Sigma-Aldrich). The following secondary antibodies were used: goat anti-rabbit IgG AlexaFluor 647 (1:400, A-21244, RRID: AB_2535812; Thermo Fisher Scientific), goat anti-mouse IgG AlexaFluor 568 (1:400, A-11004, RRID: AB_2534072; Thermo Fisher Scientific), GFP-Booster ATTO488 (1:100, gba488, RRID: AB_2631386; ChromoTek). Brightness/contrast was scaled identically within each channel for quantitative immunofluorescence experiment shown (Fig. 1B-D) and scaled individually for immunofluorescence on turbulent spindles (Fig. 1E-F).

### Microscopy

For in vivo live imaging, cells were plated either onto #1.5 glass-bottom 35 mm dishes with 23 mm well coated with poly-D-lysine (FD35PDL, World Precision Instruments, Inc.) for confinement experiments (Fig. 2, 3, and S3) or onto #1.5 glass-bottom 35 mm dishes coated with poly-D-lysine (P35G-1.5-20-C, MatTek Life Sciences) (Fig. 3B, 4, and S2A). Cells were imaged in a humidified stage-top incubator maintained at 37 °C and 5% CO_2_ (Tokai Hit). Fixed and live cells were imaged on a spinning disk (CSU-X1, Yokogawa) confocal inverted microscope (Eclipse Ti-E, Nikon Instruments) with the following components: Di01-T405/488/568/647 head dichroic (Semrock); 405 nm (100 mW), 488 nm (150 mW), 561 nm (100 mW), and 642 nm (100 mW) diode lasers; ET455/50M, ET525/36M, ET600/50M, and ET690/50M emission filters (Chroma Technology); and a Zyla 4.2 sCMOS camera (Andor Technology). Images were acquired with a 100x 1.45 Ph3 oil objective (Nikon Instruments) using Micromanager 2.0.0. FRAP experiments (Fig. 4) were performed on an OMX-SR inverted microscope (GE Healthcare) with the following components: three PCO Edge 5.5 sCMOS cameras; an environmental chamber maintained at 37 °C and 5% CO_2_ (GE Healthcare); and a Plan ApoN 60x 1.42 oil objective.

For *in vitro* imaging (Fig. 5), images were acquired using a custom-built, multicolor TIRF microscope with a Nikon inverted Ti-E microscope body, a 100X, 1.49 N.A. oil-immersion objective (Nikon), a Perfect Focus System, and an EMCCD camera (Andor iXon EM+, 512 × 512 pixels), yielding a 160 nm effective pixel size. mNeonGreen and Cy3 probes were excited with 488 nm and 561 nm coherent lasers respectively, with signals filtered through a notch dichroic filter and 525/40 and 585/40 bandpass filters (Semrock). The microscope and image acquisition were controlled via Micromanager 1.4.

### PDMS-based cell confinement assay (Fig. 2-3)

To confine cells, PDMS pillars 5 µm in height were attached to a 10 mm-diameter coverslip (4DCell), and were lowered onto cells using negative pressure generated by a Cobalt autonomous microfluidic pump (Elvesys). Pillars were lowered onto the cells over 2 min and maximum confinement was sustained for an additional 45 min. Since the area of confinement in the device is smaller than the area of the microscopy dish, cells were excluded from analysis if the final confined height was >5.1 µm, as assessed from z-stacks taken after confinement (0.3 µm step size), or if cells entered anaphase during confinement. Brightness/contrast for each channel was scaled on a per cell basis for each confinement experiment shown (Fig. 2B, 2E, S3).

### FRAP (Fig. 4)

Photobleaching was performed on a 2-µm-diameter circular region at one spindle pole. After determining the region of interest, a 488-nm laser was used to bleach the region. Images were acquired at 1 s intervals for a total of 90 s.

### Protein expression and purification (Fig. 5)

The constructs NuMA Cterm-mNeonGreen and NuMA Cterm-5A3-mNeonGreen were cloned into the pOmnibac backbone and transformed into *E. coli* DH10Bac competent cells, which were then plated on Bacmid selection plates containing Bluo-Gal. After 72 hours of incubation at 37°C, a white colony was selected and grown overnight in 2xYT medium. Bacmid DNA was purified using isopropanol precipitation and subsequently transfected into adherent SF9 cells at a density of 10^6^ cells/ml using FuGENE. When mNeonGreen expression, as assessed by fluorescence microscopy (Leica), indicated a transfection efficiency above 90%, P1 virus was harvested. To produce P2 virus, 50 ml of SF9 cells in suspension were infected with 2 ml of P1 virus and incubated for 72 hours. The cells were then centrifuged at 4,000 g for 5 minutes at 4°C, and the supernatant containing P2 virus was collected. For protein expression, 1-liter suspension of SF9 cells at 10^6^ cells/ml density was infected with 1% (v/v) P2 virus and cultured at 27°C for 72 hours. The cells were pelleted by centrifugation at 4,000 x g for 10 minutes, washed with ice-cold PBS, and flash-frozen for subsequent purification.

For protein purification, the cell pellets were lysed in lysis buffer containing 50 mM HEPES (pH 7.4), 1 M KCl, 1 mM EGTA, and 10% glycerol supplemented by 1 mM DTT, 2 mM PMSF, and two cOmplete protease inhibitor tablets (Roche) per liter of cells using a Dounce homogenizer. The lysate was clarified by centrifuging at 65,000 x g for 30 minutes, after which the supernatant was incubated with IgG Sepharose beads (Cytiva) for 1 hour at 4°C on a rotator. The beads were loaded onto a gravity-flow column and sequentially washed with lysis and then wash buffers including 50 mM HEPES (pH 7.4), 300 mM KCl, 1 mM EGTA, and 10% glycerol. To release proteins, the beads were incubated with 0.1 mg/ml TEV protease at 4°C overnight. Proteins were then eluted through gravity flow and concentrated using 30kD molecular weight cutoff (MWCO) concentrators (Amicon). Protein concentrations were measured by absorbance at 280 nm with a NanoDrop 1000 (Thermo Fisher). Proteins were loaded onto a Superdex 200 Increase 10/300 GL (Cytiva) column for size exclusion chromatography in 50 mM HEPES (pH 7.4), 150 mM KCl, 1 mM EGTA, and 10% glycerol.

### Mass photometry (Fig. 5)

High-precision coverslips (Azer Scientific) were cleaned by ultrasonically washing them three times with isopropanol followed by water for 2 minutes per cycle, then air-dried.

Gaskets were also rinsed three times in alternating water and isopropanol, dried, and placed on a cleaned coverslip. To set autofocus, 14 µL of mass photometry buffer (50 mM HEPES, pH 7.4, 150 mM KCl, 1 mM EGTA, and 10% glycerol) was added to a well in the gasket. The protein sample was prepared at a concentration of 20 nM in the same buffer and introduced into the autofocused well. Protein contrast counts were obtained using a TwoMP mass photometer (Refeyn 2) with at least two replicates. The mass photometer was calibrated with a protein mixture of conalbumin, aldolase, and thyroglobulin. Mass photometry profiles were analyzed with DiscoverMP software (Refeyn), which applied skewed Gaussian fits to the data, providing mean, standard deviation, and relative proportions for each peak.

### Flow chamber preparation (Fig. 5)

PEG-coated coverslips were prepared to minimize nonspecific protein binding. Coverslips were cleaned by ultrasonication in water, acetone, and water, then incubated in 1 M KOH for hydroxylation. After rinsing, coverslips were aminosilanized, rinsed, and treated with biotin-PEG in NaHCO_3_ buffer (pH 7.4) overnight at 4°C. These coverslips were then attached to glass slides with double-sided tape to prepare flow chambers.

### Microtubule polymerization assay (Fig. 5)

To prepare biotinylated microtubules, unlabeled, Cy3 labeled, and biotin-labeled pig brain tubulin were mixed in BRB80 buffer and combined with polymerization buffer containing GTP and DMSO, followed by incubation at 37°C. After adding Taxol, the microtubules were pelleted and resuspended in BRB80 with Taxol and DTT.

### Microtubule binding and bundling assays (Fig. 5)

To immobilize biotinylated microtubules, PEG-biotin coated flow chambers were treated with 1 mg/ml streptavidin for 3 minutes, then rinsed with MB buffer (30 mM HEPES, pH 7.4, 5 mM MgSO_4_, 1 mM EGTA, 1 mM DTT, and 10 µM Taxol). Biotinylated microtubules were introduced into the chamber, incubated for an additional 3 minutes, and then washed again with MB buffer. Different concentrations of NuMA proteins were then prepared in MBC buffer supplemented with and flowed into the chambers for imaging (Fig. 5D-E).

For the microtubule bundling assay (Fig. 5F-G), we used biotin free Cy3 labeled microtubules while keeping them near the glass surface using depletion forces generated by methylcellulose in the PEG-biotin coated flow chamber. Microtubules were mixed with 10 nM NuMA-Cterm-5A-3 or 100 nM NuMA-Cterm and incubated for 5 min at room temperature. The chamber was washed with MBC buffer (30 mM HEPES, pH 7.4, 5 mM MgSO₄, 1 mM EGTA, 0.2% methylcellulose, 1 mM DTT, 10 µM Taxol) and the mixture was then diluted to 10 µl into MBC buffer supplemented with 0.1 mg/mL glucose oxidase, 0.2 mg/mL catalase, 0.8% D-glucose and flowed into the chamber for imaging.

### Image analysis

#### Quantification of NuMA and dynein levels (Fig. 1B-D)

To compare NuMA and dynein levels in control WT RPE1 cells versus cells overexpressing NuMA-FL, NuMA-SpM, or NuMA-HCC, we quantified NuMA and p150 intensity ((integrated density – (area x mean background fluorescence))/area) at poles (2.03 µm x 2.03 µm oval selection) and spindle body (variably sized rotated rectangle selection between two poles) at the center z-slice using FIJI (ImageJ version 2.14.0/1.54f). For plots in Figure 1C and 1D, values were normalized to the mean value for WT RPE1 (control) cells.

#### Analysis for PDMS-based cell confinement assay (Fig. 2-3)

Spindle length and height were measured manually using the line selection tool in FIJI. Spindle pole fracture was defined as appearance of two distinct minus-end clusters at one pole, specifically assessed from z-stacks taken 45 minutes after confinement (Fig. 2C, 2F, 3C, and 3E). Time of spindle fracture is defined as first timepoint where two distinct minus-end clusters appear in timelapses where it could be identified.

#### Quantification of NuMA-GFP expression levels (Fig. 3D and 3F)

To assess expression level of NuMA-GFP constructs, we averaged GFP intensity (integrated density) of both poles (2.03 µm x 2.03 µm oval selection) at the center z-slice from z-stacks taken before confinement using FIJI.

#### FRAP analysis (Fig. 4)

The mean fluorescence intensity value in the bleached area, unbleached area at opposite spindle pole, and background area outside the cell were measured using a custom macro in FIJI. The signal was corrected for background and photobleaching, then normalized 0 to 1, with 0 corresponding to the lowest signal and 1 to the highest signal after photobleaching. Recovery measurements were quantified by fitting normalized fluorescence intensities of bleached areas to an exponential using GraphPad Prism. The halftimes (overall, fast, and slow) were calculated from these double exponential fits (FL, R^2^=0.82; SpM, R^2^=0.75; SpM-Bonsai, R^2^=0.82; SpM-5A-3, R^2^=0.67; Cterm-5A-3, R^2^=0.75).

#### Microtubule binding and bundling analysis (Fig. 5)

Microtubule binding images of NuMA and microtubule bundling images were analyzed in ImageJ. For the microtubule binding assay, the intensity of NuMA bound to microtubules was quantified after background subtraction using a custom MATLAB code called MTIMBS^66^, available at https://github.com/Yildiz-Lab/MTIMBS/tree/main. For the microtubule bundling assay, the intensity of microtubules was quantified after background subtraction for each condition using MTIMBS.

### Statistical analysis

Details of statistical tests and sample sizes (number of cells and number of independent experiments) are provided in figure legends. Fisher’s exact tests were performed to compare categorical datasets. Two-sided two-sample t-tests or student’s t-tests (indicated in figure legends) were performed to compare continuous datasets, based on the assumption that NuMA and dynein levels are approximately normally distributed. We used p < 0.05 as the threshold for statistical significance.

## SUPPLEMENTARY FIGURE LEGENDS

**Figure S1.**
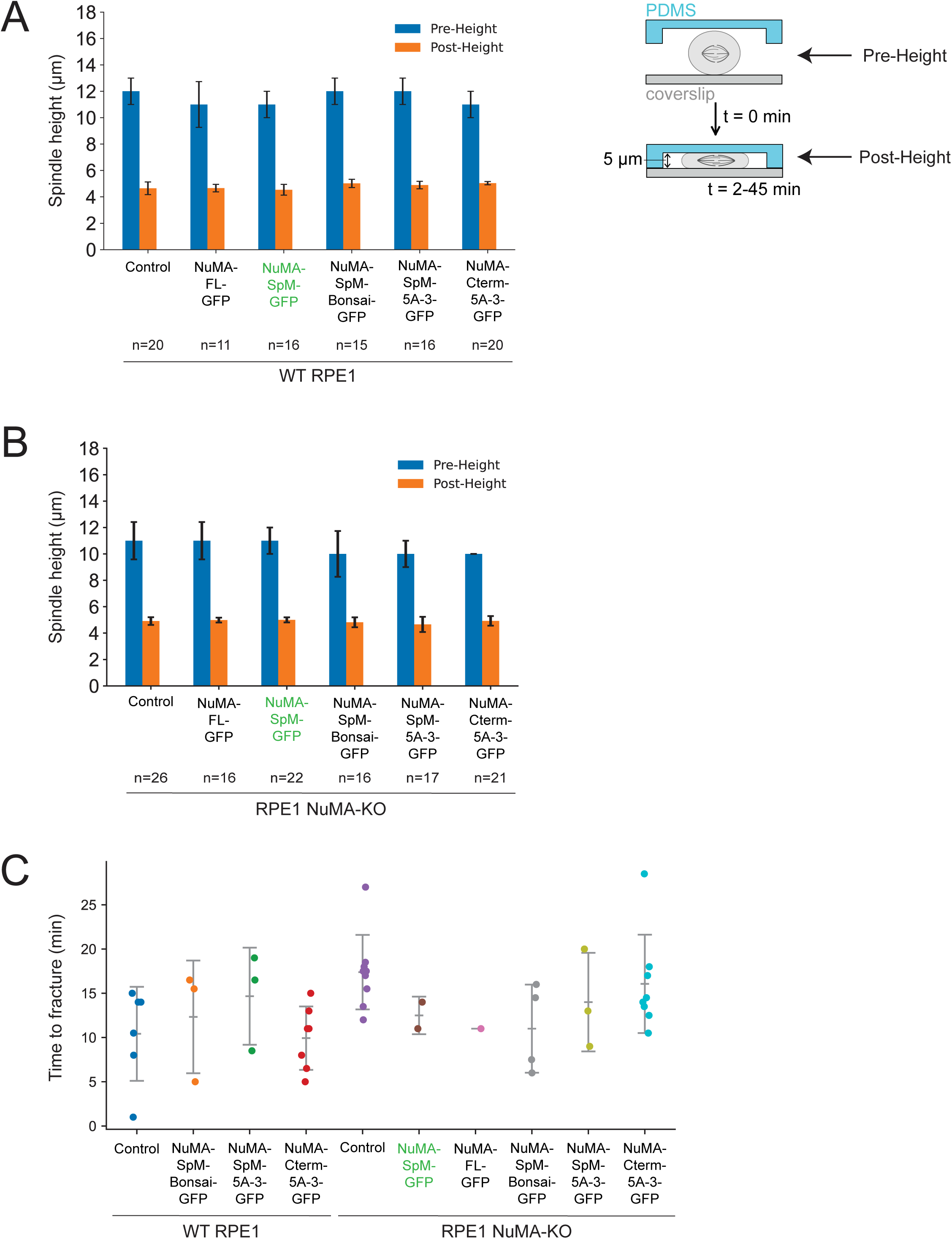
Spindle heights before and after confinement and time of spindle fracture during confinement. **(A)** Spindle heights (left panel) of WT RPE1 and RPE1 expressing NuMA-SpM, NuMA-HCC, NuMA-FL, NuMA-SpM-Bonsai, NuMA-SpM-5A-3, or NuMA-Cterm-5A-3 before and after cell confinement (right panel), as measured through confocal z-stacks. Differences between pre-heights and differences between post-heights are all not significant; two-sample t-test. Mean ± S.D. **(B)** Spindle heights of RPE1 NuMA-KO cells expressing NuMA-SpM, NuMA-HCC, NuMA-FL, NuMA-SpM-Bonsai, NuMA-SpM-5A-3, NuMA-Cterm-5A-3, or nothing (control) before and after cell confinement, as measured through confocal z-stacks. Differences between pre-heights and differences between post-heights are all not significant; two-sample t-test. Mean ± S.D. **(C)** Time of spindle fracture across all confinement conditions, which is defined as the first timepoint where the spindle pole fracture appears. n=6, 3, 3, 7, 9, 2, 1, 4, 3, 8. Differences between mean time to fracture are all not significant or not testable (for column with n=1); two-sample t-test. Mean ± S.D.

**Figure S2.**
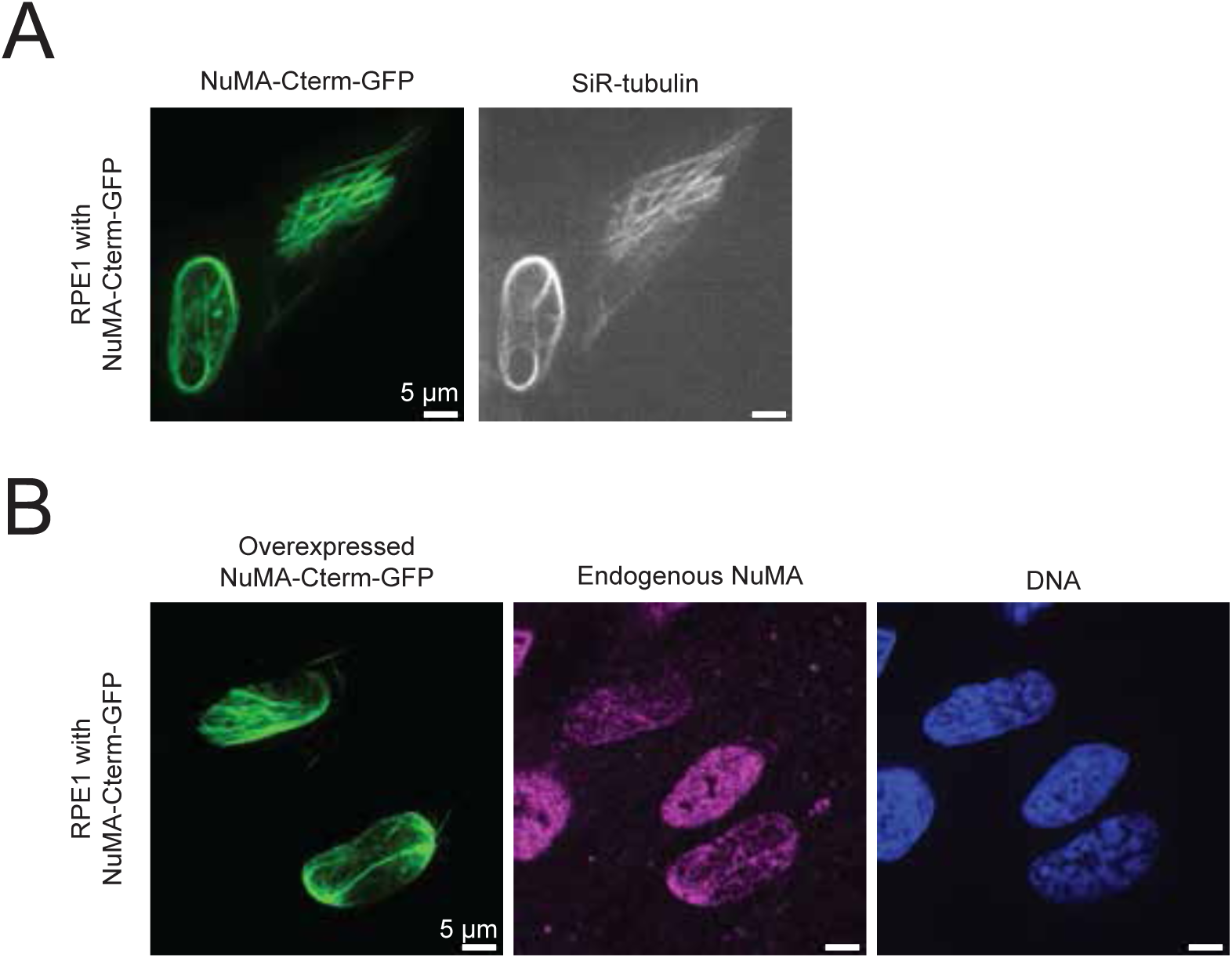
NuMA-Cterm-GFP interphase nuclear filaments. **(A)** Representative confocal images of RPE1 cells expressing NuMA-Cterm-GFP (green) after nocodazole treatment (5 μM nocodazole for 15 min), stained with SiR-tubulin (gray). NuMA-Cterm-GFP filaments persist upon nocodazole treatment and microtubules that co-localize with NuMA-Cterm-GFP also do not depolymerize. **(B)** Representative confocal images of immunofluorescence staining of RPE1 cells expressing NuMA-Cterm-GFP. Cells were stained for GFP (overexpressed NuMA), endogenous NuMA, and DNA.

**Figure S3.**
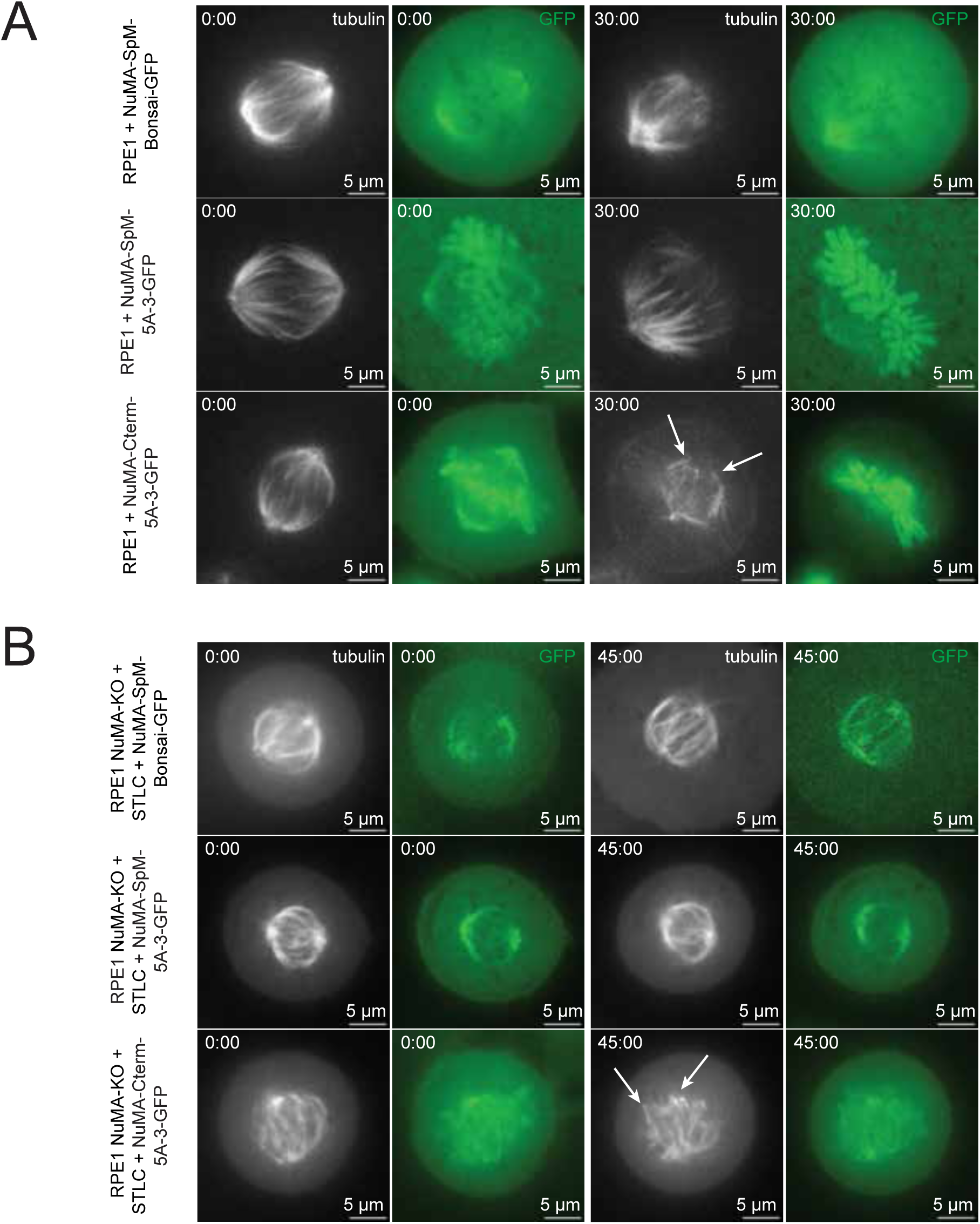
Cell confinement with combination NuMA mutants. **(A - B)** Representative confocal images taken before confinement and after significant confinement time of RPE1 (A) or RPE1 NuMA-KO + STLC (B) cells expressing NuMA-SpM-Bonsai-GFP, NuMA-SpM-5A-3-GFP, or NuMA-Cterm-5A-3-GFP. Cells were stained with either SiR-tubulin (RPE1) or with β-tubulin-HaloTag stained with Janelia Fluor 549 dye (RPE1 NuMA-KO + STLC). After cell confinement at height 5 µm, spindle pole fracture occurs in some cells, marked by white arrows.

## MOVIE LEGENDS

**Movie S1. WT RPE1 spindles fracture at a moderate rate under confinement.** Timelapse confocal imaging of representative WT RPE1 control cell, stained with SiR-tubulin, under full confinement beginning at time 0:00 (min:sec). Full confinement is reached over two minutes prior to imaging, then sustained throughout imaging. Movie corresponds to still images in Figure 2B.

**Movie S2. RPE1 spindles expressing NuMA-SpM-GFP do not fracture under confinement.** Timelapse confocal imaging of representative RPE1 cell expressing NuMA-SpM-GFP (green), stained with SiR-tubulin (gray), under full confinement beginning at time 0:00 (min:sec). Full confinement is reached over two minutes prior to imaging, then sustained throughout imaging. Movie corresponds to still images in Figure 2B.

**Movie S3. RPE1 spindles expressing NuMA-FL-GFP do not fracture under confinement.** Timelapse confocal imaging of representative RPE1 cell expressing NuMA-FL-GFP (green), stained with SiR-tubulin (gray), under full confinement beginning at time 0:00 (min:sec). Full confinement is reached over two minutes prior to imaging, then sustained throughout imaging. Movie corresponds to still images in Figure 2B.

**Movie S4. NuMA-FL-GFP turns over more slowly at spindle poles than other NuMA constructs.** Timelapse confocal imaging of representative RPE1 cell expressing NuMA-FL-GFP, NuMA-SpM-GFP, NuMA-SpM-Bonsai-GFP, NuMA-SpM-5A-3-GFP, or NuMA-Cterm-5A-3-GFP (gray). In this FRAP experiment, spindle poles are bleached after timepoint 0:02 (min:sec) and recovery is measured over the remaining time. Movies correspond to still images in Figure 4A.

